# Human Norovirus Neutralized by a Monoclonal Antibody Targeting the HBGA 1 Pocket

**DOI:** 10.1101/489906

**Authors:** Anna D. Koromyslova, Vasily A. Morozov, Lisa Hefele, Grant S. Hansman

**Affiliations:** Schaller Research Group at the University of Heidelberg and the DKFZ, Heidelberg, Germany; Department of Infectious Diseases, Virology, University of Heidelberg, Heidelberg, Germany; Pediatric Infectious Diseases Unit, University Children’s Hospital Mannheim, University of Heidelberg, Mannheim, Germany

## Abstract

Temporal changes in the GII.4 human norovirus capsid sequences occasionally result in the emergence of genetic variants capable of causing new epidemics. The GII.4 persistence is believed to be associated with the recognition of numerous histo-blood group antigen (HBGA) types and antigenic drift. We found that one of the earliest known GII.4 isolate (1974) and a more recent epidemic GII.4 variant (2012) had varied norovirus-specific monoclonal antibody (MAb) reactivities, yet similar HBGA binding profiles. To better understand the binding interaction of one MAb (10E9) that had varied reactivities with these GII.4 variants, we determined the X-ray crystal structure of the NSW-2012 GII.4 P domain 10E9 Fab complex. We showed that the 10E9 Fab interacted with conserved and variable residues, which could be associated with antigenic drift. Interestingly, the 10E9 Fab binding pocket partially overlapped the HBGA pocket and had direct competition for conserved HBGA binding residues (i.e., Arg345 and Tyr444). Indeed, the 10E9 MAb blocked norovirus VLPs from binding to several sources of HBGAs. Moreover, the 10E9 antibody completely abolished virus replication in the human norovirus intestinal enteroid cell culture system. Our new findings provide first direct evidence that competition for GII.4 HBGA binding residues and steric obstruction could lead to norovirus neutralization. On the other hand, the 10E9 MAb recognized residues flanking the HBGA pocket, which are often substituted as the virus evolves. This mechanism of antigenic drift likely influences herd immunity and impedes the possibility of acquiring broadly reactive HBGA-blocking antibodies.

**IMPORTANCE:** The emergence of new epidemic GII.4 variants is thought to be associated with changes in antigenicity and HBGA binding capacity. Here, we show that HBGA binding profiles remain unchanged between 1974 and 2012 GII.4 variants, whereas these variants showed varying levels of reactivities against a panel of GII.4 MAbs. We identified a MAb that bound at the HBGA pocket and blocked norovirus VLPs from binding to HBGAs and neutralized norovirus virions in the cell culture system. Raised against GII.4 2006 strain this MAb was unreactive to GII.4 1987 isolate, but was able to neutralize newer 2012 strain, which has important implications for vaccine design. Altogether, these new findings suggested that the amino acid variations surrounding HBGA pocket lead to temporal changes in antigenicity without affecting the ability of GII.4 variants to bind HBGAs, which are known co-factors for infection.

## INTRODUCTION

Human noroviruses are a major cause of outbreaks of acute gastroenteritis worldwide. The human norovirus genome contains three open reading frames (ORFs), where ORF1 encodes non-structural proteins, ORF2 encodes capsid protein, and ORF3 encodes a minor capsid protein (1). Human noroviruses are grouped into several genogroups (i.e., GI, GII, and GIV), which are subsequently sub-grouped into numerous genotypes. Norovirus capsid genes frequently evolve into genetically and antigenically variants, which have sometimes resulted in epidemics and even pandemics (2). The GII genotype 4 (GII.4) variants have dominated over the past two decades and have approximately 5% capsid amino acid divergence from previous epidemic GII.4 variants. The capsid of each variant likely accumulates advantageous mutations from the preceding variant. Antigenic drift has therefore been proposed as a major driving force in the emergence of novel GII.4 variants (3, 4).

Expression of the human norovirus ORF2 in insect cells results in the formation of virus-like particles (VLPs) that are antigenically and morphologically similar to native virions. The X-ray crystal structure of norovirus VLPs from the prototype strain (GI.1) identified two domains, termed shell (S) and protruding (P) domains (5). The S domain surrounds the viral RNA, whereas the P domain, which can be further subdivided into P1 and P2 subdomains, contains determinants for cell attachment and antigenicity.

Human noroviruses bind to histo-blood group antigens (HBGAs) and this interaction is important for infection (6-11). The X-ray crystal structures of the P domains from epidemic GII.4 variants in complex with a panel of HBGA showed that a regular set of conserved residues (i.e., Asp374, Arg345, Thr344, Tyr444, and Gly443) usually holds both the ABH- and Lewis-fucose of HBGAs (12-15). Interestingly, the HBGA binding profiles of the antigenically distinct GII.4 variants have likely remained unchanged over the past decades (13, 16). Indeed, the region immediately beneath the HBGA binding pocket showed little variation, whereas the surrounding residues displayed moderate amino acid substitutions (13).

Immunity to norovirus is still poorly understood (17), although volunteer studies have shown that protective immunity after infection may be absent or short-lived (18). Indeed, infection might not confer immunity against another genotypes or even variants within a genotype (19-21). Therefore, antigenic changes and HBGA binding capacities are likely an important driving force behind the GII.4 continued persistence and repeated occurrence in world-wide epidemics (19, 21).

A number of studies have discovered norovirus specific antibodies that can block HBGA binding and higher levels of antibodies that blocked VLP binding to HBGAs were associated with a lower risk of disease (22-26). Recently, an IgA MAb isolated from a volunteer infected with GI.1 norovirus was shown to have HBGA blocking activity and bound to surface-exposed loops in the vicinity of HBGA binding site (23). However, the ability of this antibody to neutralize native virus has not been determined. Moreover, the IgA was GI.1 specific and lacked blocking activity against other GI genotypes. Likewise, we discovered a Nanobody (VHH) that bound at the GII.10 HBGA pocket and showed convincing HBGA blocking potential, but was also genotype specific (27). Other Nanobodies were shown to inhibit norovirus binding to HBGAs by several distinct mechanisms, including allosteric inhibition and disruption of capsid integrity (27).

Recently, a system utilizing human intestinal enteroids (HIE) was reported to support replication of human norovirus (10). Moderate replication levels could be enhanced with addition of bile acids, however the replication success was still strongly dependent on the particular stool sample (28). This cell culture system highlighted the requirement of HBGAs on the surface of permissive cells and showed that HBGA blocking sera can effectively neutralize norovirus. Importantly, the HIE system was used to evaluate the neutralizing properties of 25 different GII.4 specific IgG and IgA isolated from a human donor (28). Several antibodies were able to neutralize human norovirus and were predicted to bind to the P domain, however the precise binding sites were not yet determined. Likewise, the mechanism of the neutralization and HBGA binding inhibition was not defined.

In this study, we examined antigenic drift with one of the earliest known GII.4 isolate (CHDC-1974) and two recent epidemic GII.4 variants (Saga-2006 and UNSW-2012). We found that GII.4 variants showed mixed MAb reactivities, while retaining HBGA binding. Following this result, we determined the X-ray crystal structure of a GII.4 variant-specific MAb (termed 10E9) in complex with the GII.4 P domain. Interestingly, the 10E9 MAb partially overlapped the HBGA pocket and this interaction blocked VLPs from binding to surrogate sources of HBGAs and neutralized norovirus in the HIE system. Overall, these data suggested that antibodies that compete with the GII.4 HBGA binding site could effectively neutralize norovirus, but antigenic drift likely limits broad antibody inhibition.

## MATERIALS AND METHODS

### VLP production

The CHDC-1974 (ACT76142), Saga-2006 (AB447457), and NSW-2012 (JX459908) VLPs were expressed as described (29). The VLPs were purified using CsCl equilibrium gradient ultracentrifugation at 35000 rpm for 24 h at 4°C (Beckman SW55 rotor). The integrity of the VLPs was confirmed by negative-stain electron microscopy (EM). VLPs were diluted in water, applied on carbon coated EM grids and stained with 1% uranyl acetate. The grids were examined on a Zeiss 910 electron microscope (Zeiss, Oberhofen, Germany) at 50,000-fold magnification.

### MAb reactivities

Eleven different IgG MAbs (Virostat, USA) were purified from a mouse immunized using the GII.4 Minerva-2006 VLPs (Genbank accession number AFJ04709.1). The Minerva capsid sequence had 99%, 95%, and 90% amino acid identity with Saga-2006, NSW-2012, and CHDC-1974, respectively. The antibody titers were quantified using a direct ELISA (30). Briefly, microtiter plates were coated with 5 μg/ml of GII.4 VLPs. The VLPs were detected with serially diluted MAbs and then a secondary HRP-conjugated goat anti-mouse MAb (Sigma, Germany). Absorbance was measured at 490 nm (OD_490_) and the binding cut-off was set to OD_490_ = 0.15 as previously determined (31). All experiments were performed in triplicate.

### HBGA binding assay

The binding of CHDC-1974 and NSW-2012 VLPs to porcine gastric mucin type III (PGM; containing HBGAs), saliva (A- and B-types), and synthetic HBGAs (A- and B-types) were measured using ELISA assays (32-35). For the PGM assay, 96-well plates were coated with 10 μg/ml of PGM for 4 h at room temperature (RT). For saliva assay, the saliva was first heated at 95°C for 10 min, briefly centrifuged, and then the supernatant was diluted 1:500 in PBS and added to wells overnight at 4°C. The PGM and saliva plates were washed three times with PBS containing 0.1% Tween20 (PBS-T), and blocked with 5% skim milk (SM) in PBS overnight at 4°C. The VLPs were two-fold serially diluted in PBS, added to duplicate wells, and then incubated for 1 h at 37°C. Plates were washed as before and reacted with HRP-conjugated goat α-mouse for 1 h at 37°C. Plates were washed and then developed with *o*-phenylenediamine and H_2_O_2_ in the dark for 30 min at RT. The reaction was stopped with 3 N HCl and absorbance was measured at 490 nm (OD_490_). For the synthetic HBGA binding assay, ELISA plates were coated with 15 μg/ml VLPs overnight at 4°C, washed with PBS-T and blocked with 5% SM. Synthetic HBGA conjugated with PAA-biotin (A-trisaccharide and B-trisaccharide; Glycotech) were two-fold serial diluted and added to duplicate wells for 2 h at 37°C. Plates were then incubated with streptavidin-HRP for 1 h at 37°C. Plates were developed as described above. All experiments were performed in triplicate.

### 10E9 Fab preparation

The 10E9 Fab (Fab #7 in this study) was prepared as described in a previous publication (29). Briefly, approximately 50 mg of purified 10E9 IgG was used for Fab preparation. The IgG was first reduced in 100 mM DTT (pH 7.6) for 1 h at 37°C. The reduced IgG was dialyzed in gel filtration buffer (GFB: 0.35 M NaCl and 2.5 mM Tris pH 7.4) supplemented with 20 mM HEPES (pH 7.7) for 1 h at 4°C. The IgG was alkylated in GFB supplemented with 2 mM iodoacetamide for 48 h at 4°C, and then transferred to GFB supplemented with 20 mM HEPES for 1 h at 4°C. The IgG was concentrated to 6 mg/ml and then digested with ficin using a commercial kit (Pierce, Rockford, USA). The Fab was separated from the Fc in a Protein A column and the resulting Fab was further purified by size exclusion chromatography with a Superdex 200 column (GE) and concentrated to 5 mg/ml and stored in GFB. The Fab sequence was determined using a commercial service (ImmunoPricise Antibodies, Canada).

### Crystallization of NSW-2012 P domain Fab complex

The NSW-2012 P domain was expressed in *E. coli* and purified as described (30, 35, 36). The NSW-2012 P domain and 10E9 Fab were mixed in a 1:1.4 molar ratio and the complex purified using size exclusion chromatography. Crystals were grown in a 1:1 mixture of the protein sample and mother liquor [0.2 M calcium acetate and 20% (w/v) PEG3350] for 6-10 days at 18°C. Prior to data collection, crystals were transferred to a cryoprotectant containing the mother liquor in 30% ethylene glycol, followed by flash freezing in liquid nitrogen.

### Data collection, structure solution, and refinement

X-ray diffraction data were collected at the European Synchrotron Radiation Facility, France at the beamline ID30A and processed with XDS (13). Structures were solved by molecular replacement in PHASER *Phaser-MR* (37) using GII.4 P domain and Fab structures (PDB: 2J4W and 4X0C) as search models. Structures were refined in multiple rounds of manual model building in COOT (38) and PHENIX (39). Structures were validated with *Procheck* (40) and *Molprobity* (41). Protein interactions were analyzed in detail using Accelrys Discovery Studio (Version 4.1) and The PyMOL Molecular Graphics System, Version 1.8 Schrödinger, LLC (42). The biologically relevant Fab-binding interface was determined using an online server (PDBePISA) and had a large surface area between the P domain and both Fab chains (heavy chain, ∼550 Å^2^ and light chain, ∼354 Å^2^). Alternative binding interfaces were located outside the CDRs and/or had small area of interaction (<250 Å^2^). Atomic coordinates and structure factors are deposited in the Protein Databank (6EWB).

### 10E9 Fab blocking assay

Blocking assays were performed as descried earlier (34). Briefly, 0.5 μg/ml of Saga-2006 and NSW-2012 VLPs were pre-treated with serially diluted 10E9 Fab for 1 h at RT and added to the PGM or saliva coated plates. The CHDC-1974 VLPs were not examined in this binding assay, since the VLPs did not bind to MAb 10E9. PBS was used as blank and untreated VLPs were used as a reference control. The OD_490_ value of untreated VLPs was set as the reference value corresponding to 100% binding. The percentage of inhibition was calculated as [1-(treated VLP mean OD_490_/mean reference OD_490_)] × 100. IC_50_ values for different inhibitors were calculated using GraphPad Prism 6.0a.

### Isothermal Titration Calorimetry measurements

Isothermal Titration Calorimetry (ITC) experiments were performed using an ITC-200 (Malvern, UK). Titrations were performed at 25°C by injecting consecutive (1-2 μl) aliquots of NSW-2012 P domain (80 µm) into 10E9 MAb (8 µM). Injections were performed until saturation was achieved. To correct for the heat of dilution, control experiments were performed with titrating the P domain into PBS. The heat associated with the control titrations was subtracted from the raw binding data prior to fitting. The data was fitted using a single set-binding model in Origin 7.0 (OriginLab, Northampton, MA). Binding sites were assumed to be identical.

### Human intestinal enteroid culture

Secretor-positive jejunal HIE culture (J2) and Noggin producing cell lines were kindly provided by M. Estes, Baylor College of Medicine, Texas, USA (10). R-sponding producing cell line and Wnt3 producing cell line were commercially obtained from Trevigen and ATCC, respectively. Wnt3a, R-spondin, and Noggin-conditioned media were produced as was reported previously (10). Wnt3 activity was tested using TCF reporter plasmid kit (Merck). Complete growth media (CMGF+), basal media (CMGF-) and differentiation media were all prepared as described in (10). HIE were grown as 3D cultures in Matrigel in CMGF+ as previously described (10). For infection experiments, HIE were trypsinized and grown as 2D monolayers in a collagen coated 96 well plates as previously reported (10). After one day the CMGF+ media was changed to differentiation media for five days. Monolayers were pretreated with 500 μM GCDCA for two days before infection experiments.

### HIE infection and inhibition

Confluent monolayers were washed once with ice cold CMGF- and incubated at 37°C for 1 h with diluted stool samples. Stool samples were kindly provided by M. Estes, H. Schroten (Mannheim Children Hospital, Mannheim, Germany), and P. Schnitzler, (Heidelberg University Clinic, Heidelberg, Germany). After infection monolayers were washed twice with ice cold PBS, and incubated in differentiation media supplemented with 500 μM GCDCA or 500 μM GCDCA and 1% human bile. Samples were frozen at 0 days post infection (dpi), i.e., 1 h after virus attachment and 4 dpi. For inhibition experiments 10E9 MAb was diluted in PBS to a stated concentration, mixed with the virus in 1:1 ratio and incubated for 1 h at 37°C. After preincubation the samples were diluted with CMGF-with 500 μM GCDCA and applied on washed HIE monolayers for 1 h at 37°C. Isotype IgG (anti-myosin heavy chain antibody, Abcam) was used as a negative control. Plates were washed and incubated as above. All experiments were performed three times with technical duplicates or triplicates.

### RNA extraction and RT-qPCR

Viral RNA was extracted at 0 and 4 dpi using RNAeasy mini kit (Quagen) or phenol/chlorophorm extraction with Ribozol according to manufacturer instructions. Genome copy levels were measured using qScript XLT One-step RT-qPCR ToughMix reagent with ROX (Quanta Bioscineces) using COG2R/ QNIF2D primer pair, and probe QNIFS as described previously (10, 28). A standard curve based on human norovirus RNA transcript was used to quantitate viral genome equivalents. Results were analyzed using Microsoft Excel and GraphPad Prism (6.0a). Statistical analysis was performed using one-way ANOVA test. Differences were considered significant when P ≤ 0.05. IC_50_ values were calculated using GraphPad Prism.

## RESULTS

### Antibody reactivities against VLPs

In order to analyse temporal changes in antigenicity, we examined the reactivities of the CHDC-1974, Saga-2006, and NSW-2012 VLPs (Fig. 1A) with 11 different GII.4 norovirus MAbs (termed MAbs #1 to #11) that were produced in a mouse immunized with the GII.4 Minerva-2006 isolate. We found that 11 MAbs reacted against the Saga-2006 VLPs (Fig. 1B). This result was not surprising, since the Minerva-2006 and Saga-2006 capsid sequences had only two amino acid differences. On the other hand, the CHDC-1974 VLPs only reacted with 7 of 11 MAbs, while the NSW-2012 VLPs reacted with 9 of 11 MAbs (Figs. 1C and 1D). Both CHDC-1974 and NSW-2012 VLPs failed to react to MAb #10. In addition, CHDC-1974 VLPs failed to bind MAbs #2, #7 (i.e., 10E9 MAb), and #8, while NSW-2012 VLPs failed to react with MAb #1. This result was likely due to lower amino acid identities of NSW-2012 and CHDC-1974 capsids, which had 26 and 50 amino acid substitutions compared to the Minerva-2006 capsid, respectively. Moreover, these results indicated that several MAbs recognised equivalent epitopes on CHDC-1974 and NSW-2012 VLPs, while other MAb binding epitopes were likely unique and associated with antigenic drift.

**Figure 1.**
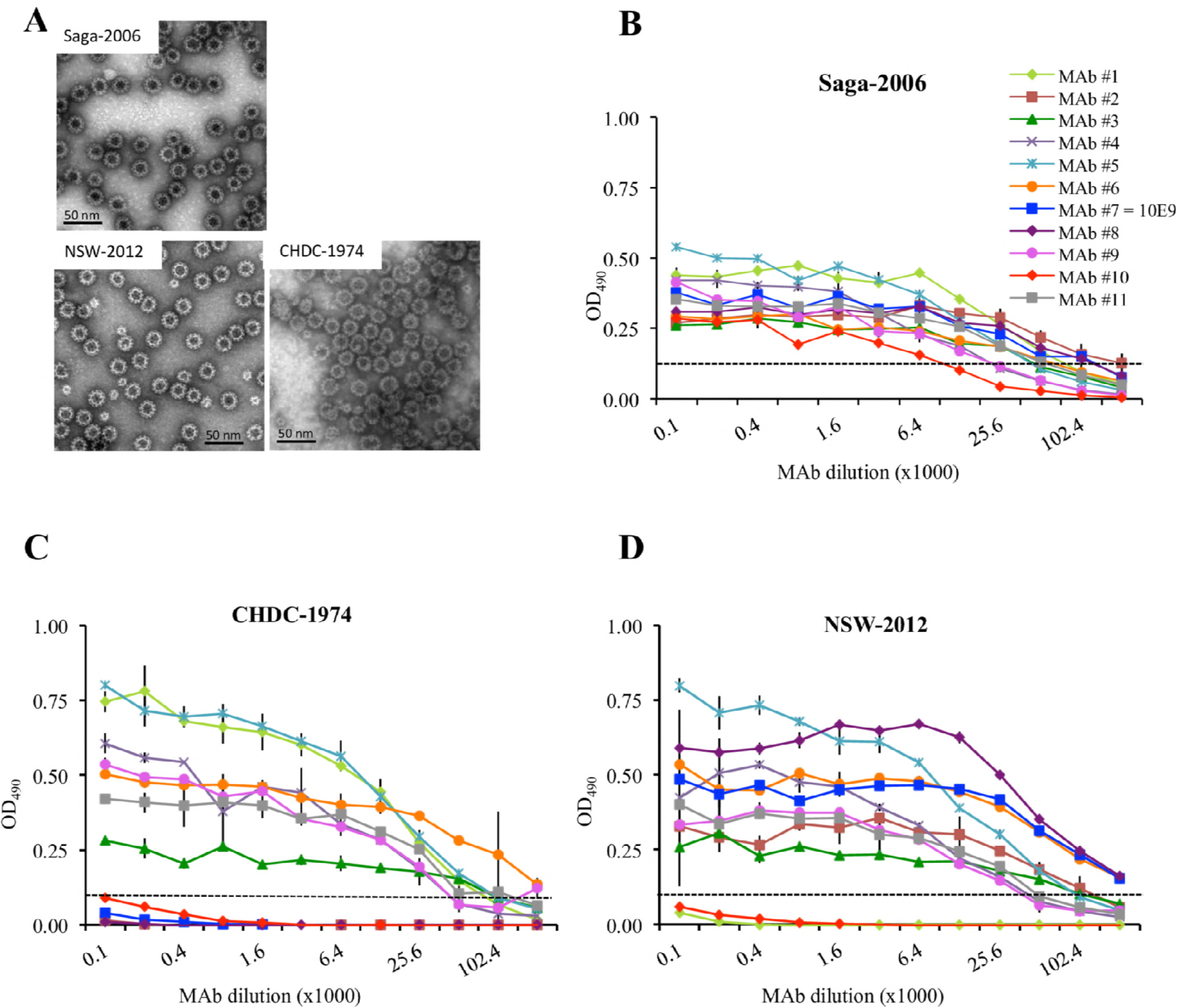
GII.4 VLP binding interactions with norovirus specific MAbs. An antigen ELISA was used to determine the cross-reactivities of the CHDC-1974 and NSW-2012 VLPs with 11 different MAbs. MAbs were two-fold serially diluted and the binding cut-off was set to OD_490_ = 0.15 (35). (A) Representative negative stain EM images of Saga-2006, NSW-2012, and CHDC-1974 VLPs. (B) The Saga-2006 VLPs reacted against all MAbs. (C) The CHDC-1974 VLPs reacted with 7 of 11 MAbs and failed to react with MAbs #2, #7, #8, and #10. (D) The NSW-2012 VLPs reacted with 9 of 11 MAbs and failed to react with MAbs #1 and #10. The MAb #7 was also termed 10E9 MAbs in this study. All experiments were performed in triplicates; standard deviation is shown with error bars.

### VLP binding interactions with HBGAs

Our previous structural study of Farmington Hills-2004, Saga-2006, and NSW-2012 GII.4 variants confirmed that these noroviruses were capable of binding numerous HBGA types, despite having varied residues interacting with the terminal HBGA saccharides (13). To compare the HBGA binding interactions of one of the earliest known GII.4 strain (CHDC-1974) with the prevalent NSW-2012 variant, we performed different HBGA binding assays using the CHDC-1974 and NSW-2012 GII.4 VLPs (Fig. 2). We found that both the CHDC-1974 and NSW-2012 VLPs bound to pig gastric mucin (PGM) in a dose-dependent manner. The CHDC-1974 VLPs bound to PGM at 0.47 µg/ml, while the NSW-2012 VLPs bound at 0.31 µg/ml (Fig. 2A). Similarly, the CHDC-1974 and NSW-2012 VLPs bound A- and B-type saliva at equivalent concentrations, approximately 0.63 µg/ml (Figs. 2B and 2C). In contrast, VLP binding to synthetic HBGAs was slightly lower, where CHDC-1974 and NSW-2012 VLPs bound to synthetic A- and B-trisaccharide at 6.25 µg/ml and 15 µg/ml, respectively (Figs. 2D and 2E). These results indicated that the CHDC-1974 and NSW-2012 VLPs were both capable of binding the A- and B-type HBGAs. Together with our previous structural data, these results suggested that these GII.4 variants maintained similar HBGA binding interactions.

**Figure 2.**
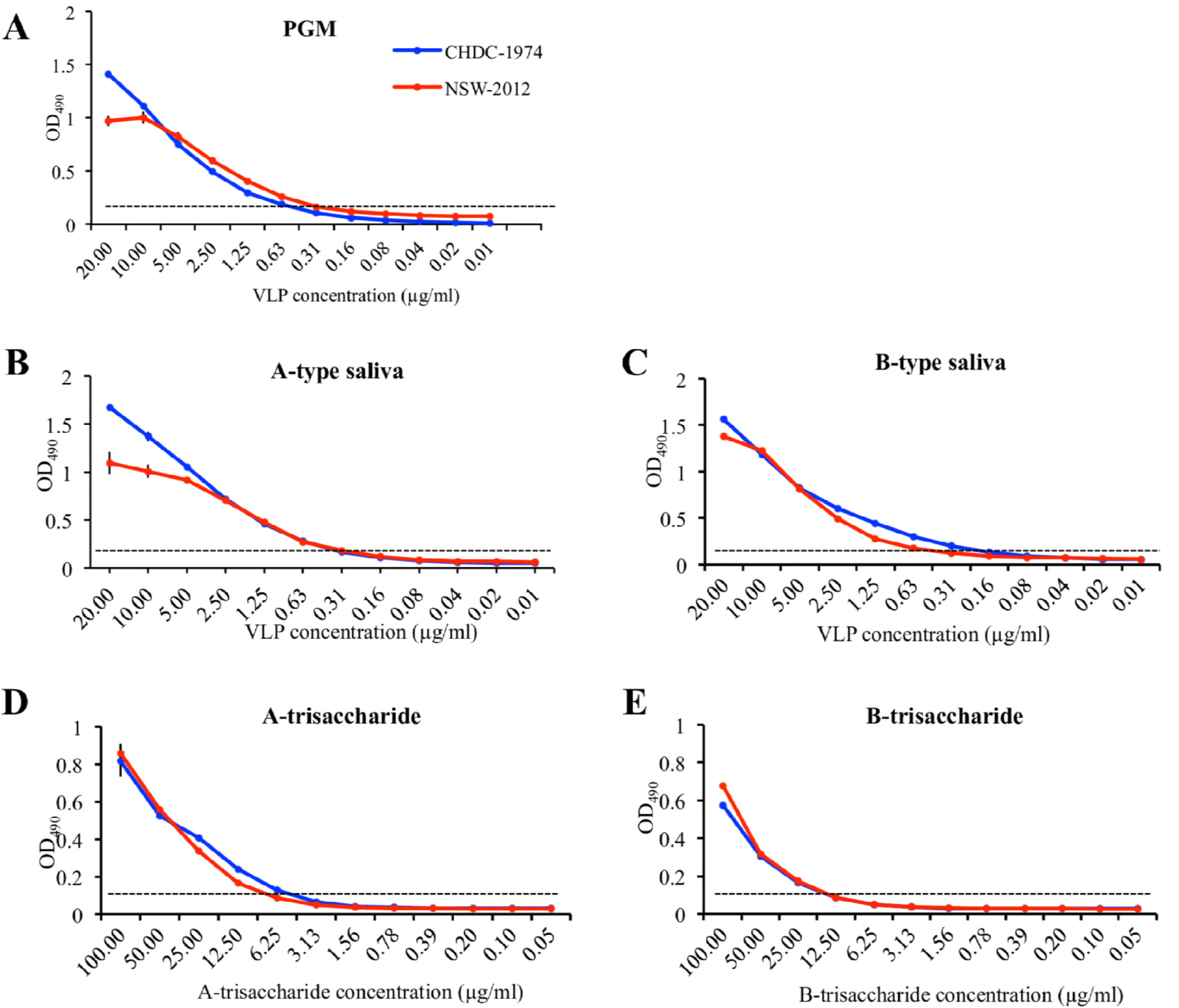
VLP binding interaction with PGM, saliva, and synthetic HBGAs. (A) The binding of CHDC-1974 and NSW-2012 VLP binding to PGM was measured using an ELISA (34). VLPs were two-fold serially diluted and the binding cut-off was set to OD_490_ = 0.15 (35). The CHDC-1974 VLPs bound to PGM with at a minimum concentration of 0.47 µg/ml, while the NSW-2012 VLPs bound at 0.31 µg/ml. (B) The CHDC-1974 and NSW-2012 VLPs bound to A-type saliva at 0.31 µg/ml. (C) The CHDC-1974 and NSW-2012 VLPs bound to B-type saliva at 0.22 µg/ml and 0.47 µg/ml, respectively. (D) The CHDC-1974 and NSW-2012 VLPs bound at 6.25 µg/ml synthetic A-trisaccharide. (E) The CHDC-1974 and NSW-2012 VLPs bound at 15 µg/ml synthetic B-trisaccharide. All experiments were performed in triplicates; standard deviation is shown with error bars.

An amino acid alignment of the CHDC-1974 P domain with the 2004, 2006, and 2012 GII.4 variants revealed that most substitutions occurred in the P2 subdomain (Fig. 3). A single amino acid insertion located at position 394 distinguished the older (i.e., 1974 strain) and newer GII.4 sequences (i.e., 2004 and onwards). The P domain residues interacting with the ABH-fucose moiety of HBGAs have for the most part remained highly conserved among these GII.4 variants (Fig. 3).

**Figure 3.**
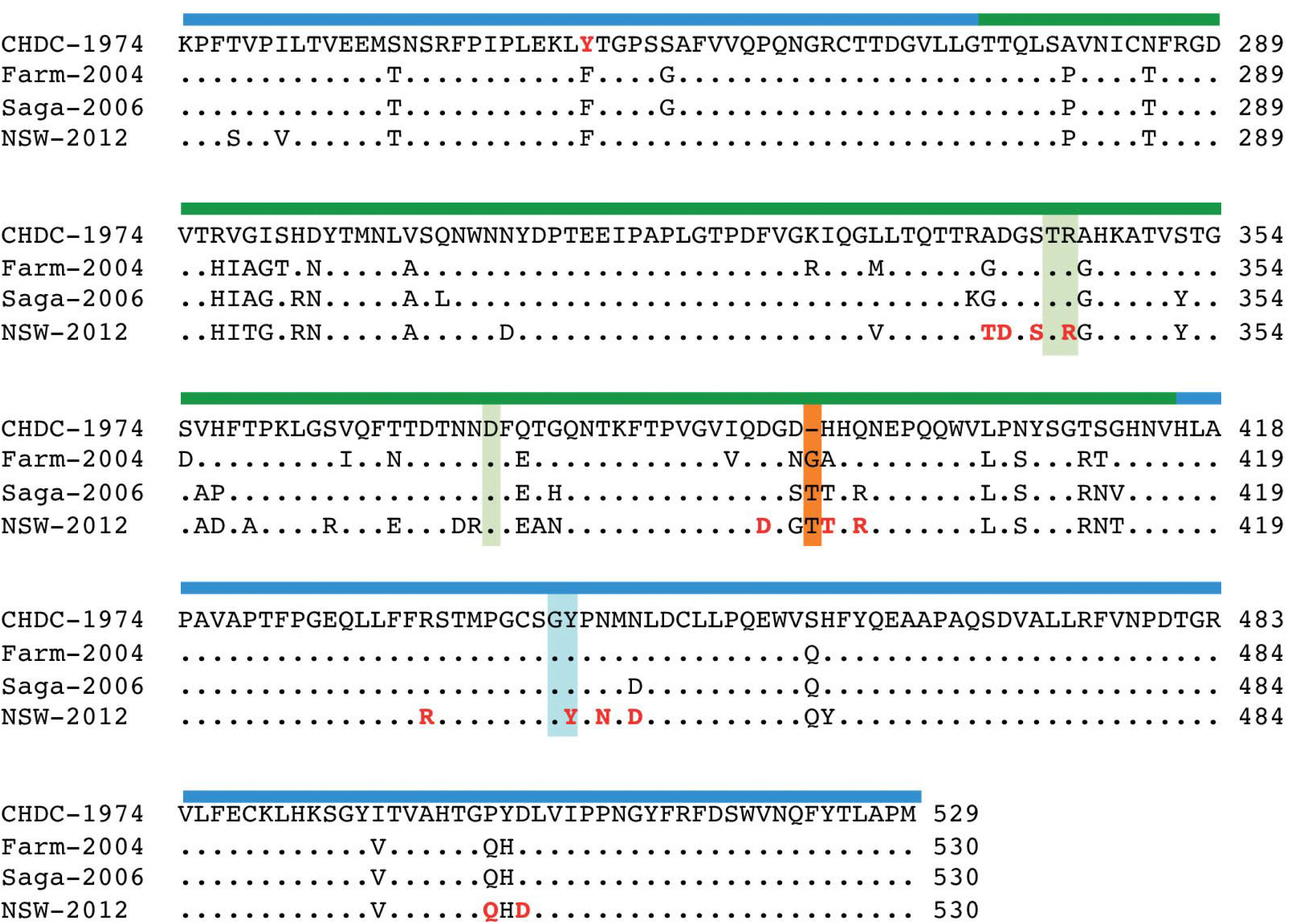
Sequence alignment of GII.4 variants. The P domain amino acid sequences of GII.4 variants CHDC-1974, Farmington Hills-2004 (Farm-2004: JQ478408), Saga-2006, and NSW-2012 were aligned. The P1 subdomain (blue bar) was mostly conserved, whereas the P2 subdomain (green bar) was more variable. The regular set of GII.4 amino acids interacting with fucose of HBGAs was shaded in light green (chain A) and light blue (chain B). Compared to the CHDC-1974 sequence, one amino acid insertion (orange shade) was found in 2004 and remained in 2006 and 2012 variants. The NSW-2012 P domain amino acids interacting with 10E9 Fab were marked in bold red.

### Structure of NSW-2012 P domain 10E9 Fab complex

The ELISA data showed that MAb #7 (also termed 10E9 MAb) was reactive against Saga-2006 and NSW-2012, but unreactive against the CHDC-1974 VLPs. This result suggested that 10E9 MAb bound at a partially variable region on the capsid that was conserved in both Saga-2006 and NSW-2012, but different in CHDC-1974. To better understand these 10E9 MAb reactivities, we determined the X-ray crystal structure of the NSW-2012 GII.4 P domain 10E9 Fab complex. A single NSW-2012 P domain 10E9 Fab crystal diffracted to 2.78 Å resolution and contained two P dimers and four Fab molecules in space group P22_1_2_1_. Data statistics are provided in Table 1. The 10E9 Fab bound at the upper-side of the P domain (Fig. 4A). The overall structure of the P domain in the P domain-Fab complex structure was reminiscent of the unbound P domain, except for some minor loop movements (chain A: residues 338-342; and chain B: residues 389-399) (13).

**Table 1.** Data collection and refinement statistics for NSW-2012 P domain 10E9 Fab structure

| Parameter | 6EWB |
| --- | --- |
| Data collection |  |
| Space group | $P22_12_1$ |
| Cell dimensions |  |
| $a, b, c$ (Å) | 106.84 111.62 288.25 |
| $\alpha, \beta, \gamma$ (°) | 90 90 90 |
| Resolution range (Å) | 48.76-2.78 (2.88-2.78)* |
| $R_{\text{merge}}$ | 15.6 (97.5)* |
| $I/\sigma I$ | 17.97 (2.7)* |
| Completeness (%) | 98.80 (92.9)* |
| Redundancy | 13.9 (12.8)* |
| CC1/2 | 84.0 |
| Refinement |  |
| Resolution range (Å) | 48.76-2.78 |
| No. of reflections | 86664 |
| $R_{\text{work}}/R_{\text{free}}$ | 18.59/22.19 |
| No. of atoms | 21709 |
| Protein | 21701 |
| Ligand/ion | 8 |
| Water | 0 |
| Average $B$ factors (Å<sup>2</sup>) | |
| Protein | 59.10 |
| Ligand/ion | 65.50 |
| Water | 0 |
| RMSD |  |
| Bond lengths (Å) | 0.005 |
| Bond angles (°) | 0.95 |
| Ramachandran favored (%) | 97 |
| Ramachandran outliers (%) | 0.14 |
| MolProbity score | 1.3 |

Each data set was collected from single crystals, respectively.
\*Values in parentheses are for highest-resolution shell.

**Figure 4.**
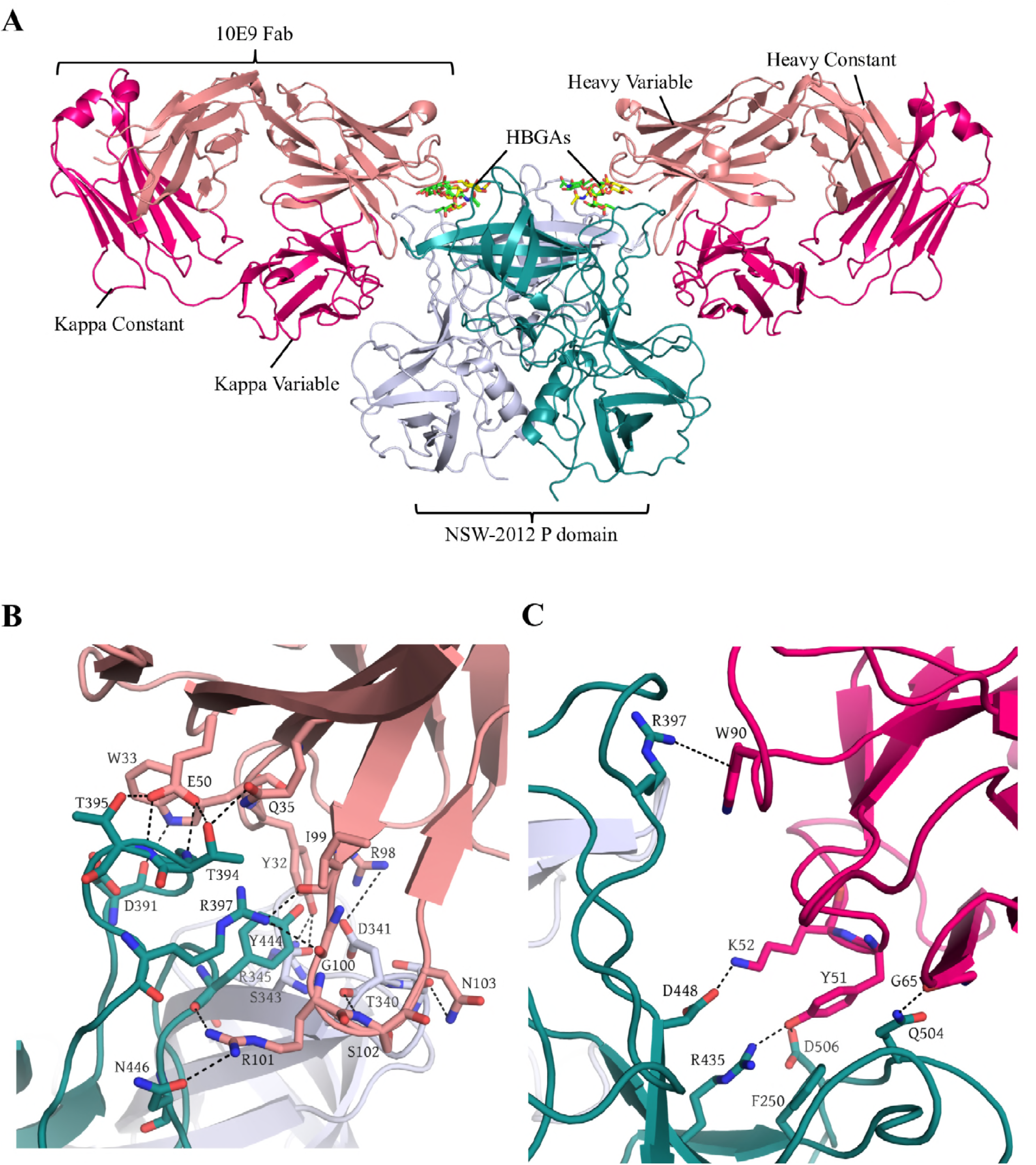
X-ray crystal structure of NSW-2012 P domain 10E9 Fab complex. The NSW-2012 P domain 10E9 Fab complex was crystallized in an orthorhombic unit cell. Lewis X (PDB ID 4X0C: yellow sticks) and A-trisaccharide (4WZT: green sticks) were superpositioned on the structure to show the HBGA binding pocket. (A) The 10E9 Fab bound to the surface exposed loops on the top of the P1 subdomain (P domain chain A: blue-white and P domain chain B: deep teal; Fab light chain: hot-pink and Fab heavy chain: salmon). (B) A network of direct 15 hydrogen bonds was formed between the Fab heavy chain and P domain monomers, i.e., P domain chain A (Thr394, Thr395, Asp391, Arg397, Tyr444, Asn446, and Asp448) and P domain chain B (Thr340, Asp341, Ser343, and Arg345). Three electrostatic interactions were also observed (Fab Glu50 and P domain chain A Arg397; and Fab Arg98 and P domain chain B Asp341). (C) The Fab light chain was involved in four direct hydrogen bonds with the P domain chain A (Gln504, Asp506, and Arg435). Additionally, the Fab light chain was coordinated with two hydrophobic interactions between Fab Tyr51 and P domain Phe250 (chain A) and between Fab Trp90 and uncharged part of Arg397 on the P domain. Two electrostatic interactions were also observed (Fab Trp90 and P domain chain A Arg397; Fab Lys52 and P domain chain A Asp448).

The binding interface presented an exceptional surface complementarity and involved numerous hydrogen bonds and hydrophobic interactions (Fig. 4B). The heavy chain of 10E9 Fab interacted with both P domain monomers (interface area 349Å^2^ with chain B and 239Å^2^ with chain A), whereas the light chain was only involved in monomeric interactions (interface area 543Å^2^). Interacting residues were primarily located in the heavy chain CDRH1 and CDRH3 and the light chain CDRL2 and CDRL3. Fourteen P domain residues (chain A: Asp391, Thr394, Thr395, Arg397, Arg435, Tyr444, Asn446, Asp448, Gln504, and Asp506; chain B: Thr340, Asp341, Ser343, and Arg345) formed nineteen direct hydrogen bonds with 10E9 Fab. Two P domain residues (chain A: Arg397; and chain B: Asp341) were involved in five electrostatic interactions. Two hydrophobic interactions were formed between Phe250, Arg397 (P domain chain A) and Tyr51, Trp90 (Fab light chain). A similar set of binding interactions was observed with the second Fab molecule.

An amino acid sequence alignment showed that most (13/14) of the NSW-2012 P domain-Fab binding residues were conserved in Saga-2006, whereas in the CHDC-1974 only 8 of 14 were shared (Fig. 3). These results indicated that the residues substituted in CHDC-1974 inhibited 10E9 MAb from binding to CHDC-1974 VLPs (Fig. 1C). Also, two amino substitutions in Minerva and Saga-2006 were located at residues 333 (Saga-2006 numbering) and 536. These substitutions were outside the HBGA pocket and not involved in 10E9 MAb binding, which suggested Minerva and Saga-2006 likely had comparable antigenicity, although Minerva VLPs were not available for testing.

### 10E9 Fab overlapped the HBGA pocket

The X-ray crystal structure of the NSW-2012 P domain-Fab revealed that the Fab bound nearby the HBGA pocket (Fig. 4). Superposition of the NSW-2012 P domain 10E9 Fab complex onto the NSW-2012 and Saga-2006 P domain HBGA complex structures revealed that the side-chains of the Fab residues clashed with the HBGAs (Fig. 5). These results indicated that 10E9 MAb might block norovirus from binding to HBGAs not only through a direct competition for residues in the HBGA binding site (Arg345 and Tyr444), but also by steric hindrance with HBGA pocket (Figs. 5B and 5C). A similar mechanism of inhibition was recently discovered with a GI.1 IgA MAb isolated from a volunteer infected with human norovirus (23). Comparison of these two complex structures indicated that partial MAb overlap of the GI.1 and GII.4 HBGA pocket could provide HBGA blocking capabilities (Fig. 5D).

**Figure 5.**
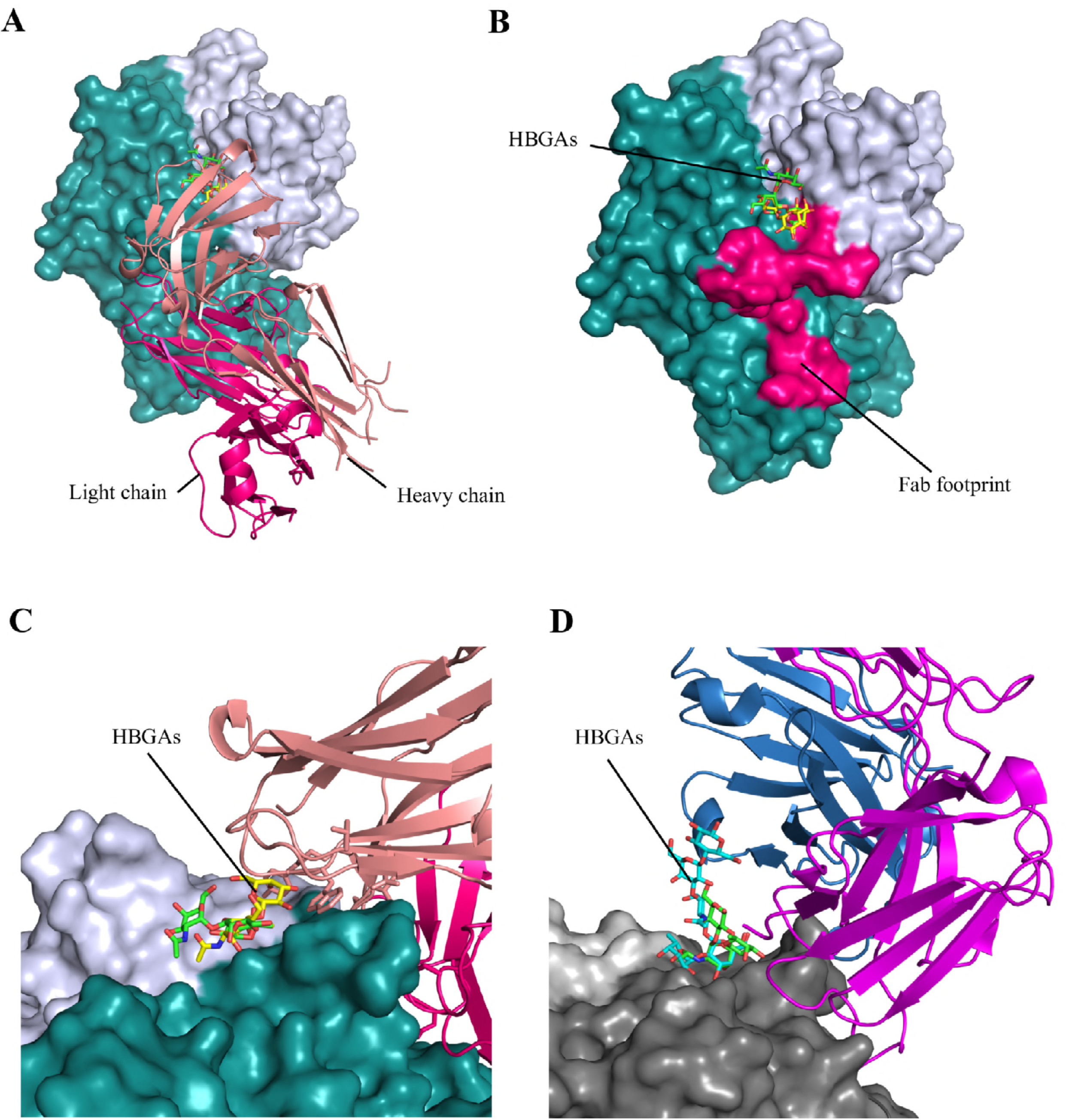
The 10E9 Fab binding epitope overlapped the GII.4 P domain HBGA binding pocket. The 10E9 Fab clashed with HBGAs and partially overlapped the HBGA binding site. The NSW-2012 P domain and 10E9 Fab was colored as in Figure 2. (A) The 10E9 Fab binding site was viewed from the top left corner of the P domain. Lewis X (yellow sticks) and A-trisaccharide (green sticks) were superpositioned on the structure to show the HBGA binding pocket. (B) The Fab footprint (hot-pink) showing the binding to two P domain monomers. Interestingly, the Fab also interacted with two P domain residues (Arg345 and Tyr444) that were involved in HBGA binding. (C) The 10E9 Fab clashed with HBGAs bound on the GII.4 P domain. (D) The GI.1 P domain Fab complex (5KW9) showing the Fab clashed with the bound HBGAs. The structure was colored accordingly: heavy chain (blue), light chain (purple), P domain chain A (light gray), P domain chain B (dark gray), H-type 1 (cyan; 2ZL6), and A-trisaccharide (green; 2ZL7).

### HBGA blocking properties of 10E9 Fab

In order to determine the HBGA blocking potential of the 10E9 MAb, we analyzed Saga-2006 and NSW-2012 VLP inhibition in the well-established HBGA surrogate neutralization assay (32-35). We found that the 10E9 Fab inhibited the Saga-2006 VLPs binding to PGM and saliva in a dose-dependent manner (Fig. 6). The 10E9 Fab IC_50_ in the PGM assay = 20 μg/ml, in the A-type saliva assay the IC_50_ = 7.9 μg/ml, and in the B-type saliva assay the IC_50_ = 6.7 μg/ml. The 10E9 Fab failed to effectively block the NSW-2012 VLPs from binding to PGM and saliva, where at the highest concentration only 35% inhibition was observed. Thus, although 10E9 recognized both Saga-2006 and NSW-2012 at comparable levels, they were distinguished in the HBGA blocking assay. The reason for this difference in inhibition was not apparent, although amino acid variations surrounding the Saga-2006 and NSW-2012 HBGA pockets (Fig. 3) might exert additional influence on the efficiency with which Saga-2006 and NSW-2012 VLPs bind to HBGAs present in PGM. Indeed, a similar kind of varied level of HBGA binding and inhibition was also observed with genetically related GII.4 noroviruses isolated from a chronically infected individual (33).

**Figure 6.**
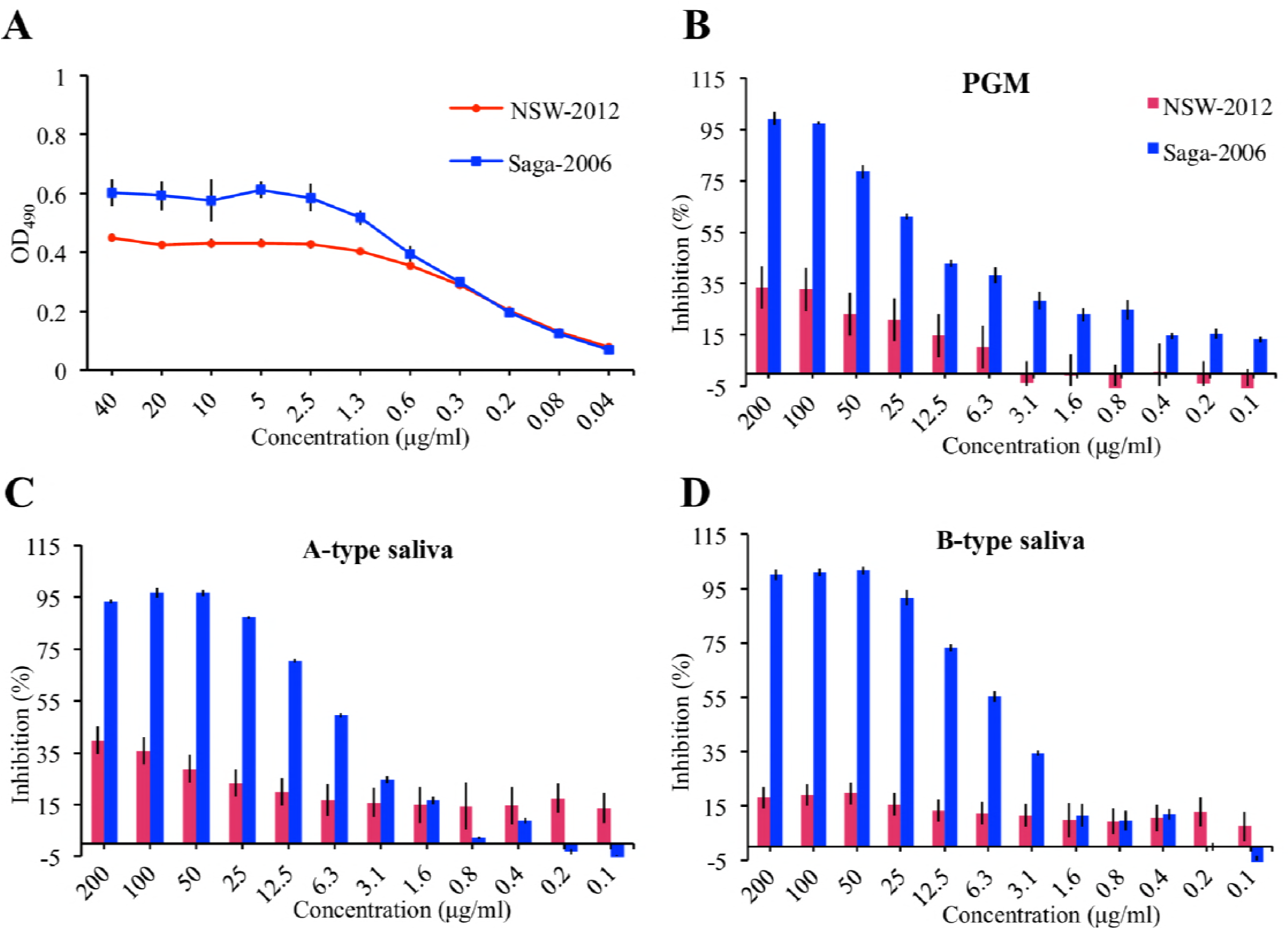
The 10E9 Fab inhibition assay. (A) The 10E9 MAb bound to Saga-2006 and NSW-2012 VLPs at comparable levels (also see Figure 1). The Saga-2006 and NSW-2012 VLPs were pretreated with serially diluted 10E9 Fab and then added to triplicate wells coated with (B) PGM, (C) A-type saliva, and (D) B-type saliva. For the Saga-2006 VLPs, the PGM assay for 10E9 Fab IC_50_ value = 20 μg/ml, for the A-type saliva assay, the 10E9 Fab IC_50_ = 7.9 μg/ml, and for the B-type saliva assay, the 10E9 Fab IC_50_ = 6.7 μg/ml. For the NSW-2012 VLPs, the 10E9 Fab IC_50_ for the PGM and saliva was not calculated, since only approximately 15-35% of inhibition was achieved at the highest Fab concentration.

### Thermodynamic properties of 10E9 IgG binding

Overall, our data indicated that the 10E9 MAb might function as GII.4 HBGA inhibitor. In order to determine the thermodynamic properties of 10E9 MAb binding to NSW-2012 P domain we analyzed the binding interaction using ITC (Fig. 7). The 10E9 IgG binding to the P domain was characterized with exothermic type of reaction with K_d_ value = 59 nM. Binding enthalpy (ΔH) of -4.3 kcal/mole and entropy change (ΔS) of 18.8 cal/mol/deg contributed almost equally to the binding affinity. Thermodynamic data from the 10E9 IgG binding to the P domain indicated that both hydrogen bonds and hydrophobic interactions were involved in binding. The 10E9 IgG affinity was equivalent to the GI.1 IgA Fab (K_d_ = 20 nM) (23). Interestingly, although MAbs have two antigen recognition regions per molecule, the stoichiometry of 10E9 IgG binding to the P domain was ∼1 (0.98). This result indicated that each antibody binds one monomer of the P domain. As the P domain forms a dimer in solution, it is likely that each P domain dimer bound two 10E9 MAb molecules.

**Figure 7.**
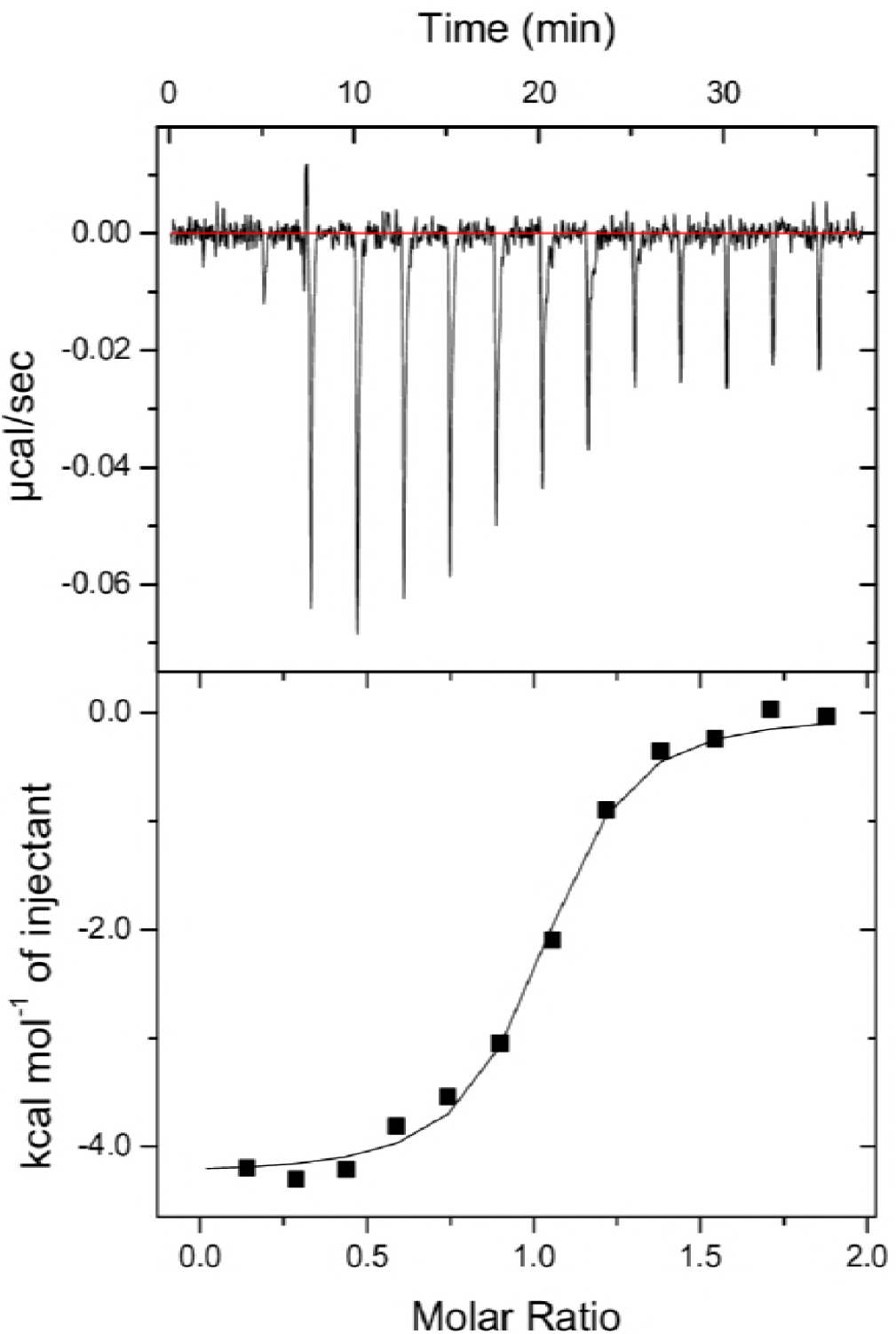
Thermodynamic properties of 10E9 IgG binding to NSW-2012 P domain. The upper panel show the heat of titrations of NSW-2012 P domain to 10E9 IgG. The bottom panel illustrates the binding isotherm. The binding isotherm was calculated using a single binding site model. The 10E9 IgG showed nanomolar affinity with Kd = 5.9E-08 M (± 2E-08 M), enthalpy ΔH = -4.3E + 03 cal/mol (±11), and entropy ΔS= 18.8 cal/mol/deg (± 2). Gibs free energy was calculated to be ΔG= - 9.9E + 03 cal/mol (± 2E-02). The binding constants corresponded to an average of three independent experiments.

### 10E9 MAb neutralization of GII.4 norovirus

The HIE system was evaluated in order to confirm the potential neutralizing properties of the 10E9 MAb with two different norovirus-positive stool samples, i.e., the previously validated GII.4 TCH12-580 (10) and another GII.4 stool sample with low, but consistent replication levels (GII.4 Mannheim 2018). Of note, we encountered difficulties in finding a stool sample with good replication levels as all ten tested samples showed limited increase in genome copies (<50 times) despite high virus load. Diluted stool samples were preincubated with the 10 μg/ml of 10E9 IgG before HIE infection. At this concentration, complete neutralization of the virus was observed for both stool samples at 4 dpi (Figs. 8A and 8B). In order to further analyze the neutralization properties, the 10E9 MAb was serially diluted and then incubated with the GII.4 TCH12-580 stool sample. We found that the 10E9 neutralized GII.4 TCH12-580 in a concentration dependent manner with IC_50_ value of 97 ng/ml (Fig. 8C). This result was different to the PGM blocking assay, which showed only ∼15% inhibition with the genetically related UNSW-2012 VLPs and was ∼200 times lower than IC_50_ (20 μg/ml) in HBGA blocking assay for GII.4 Saga-2006 VLPs (Fig. 6). Interestingly, 10E9 IgG reduced the number of input virus particles compared to untreated sample as seen by the reduction of genome copies at 0 dpi (Figs. 8 A and 8B). This result indicated that 10E9 IgG reduced the virus attachment to the cell monolayer by preventing the binding to HBGAs on the cell surface. Overall, these results confirmed that 10E9 MAb inhibited norovirus replication.

**Figure 8.**
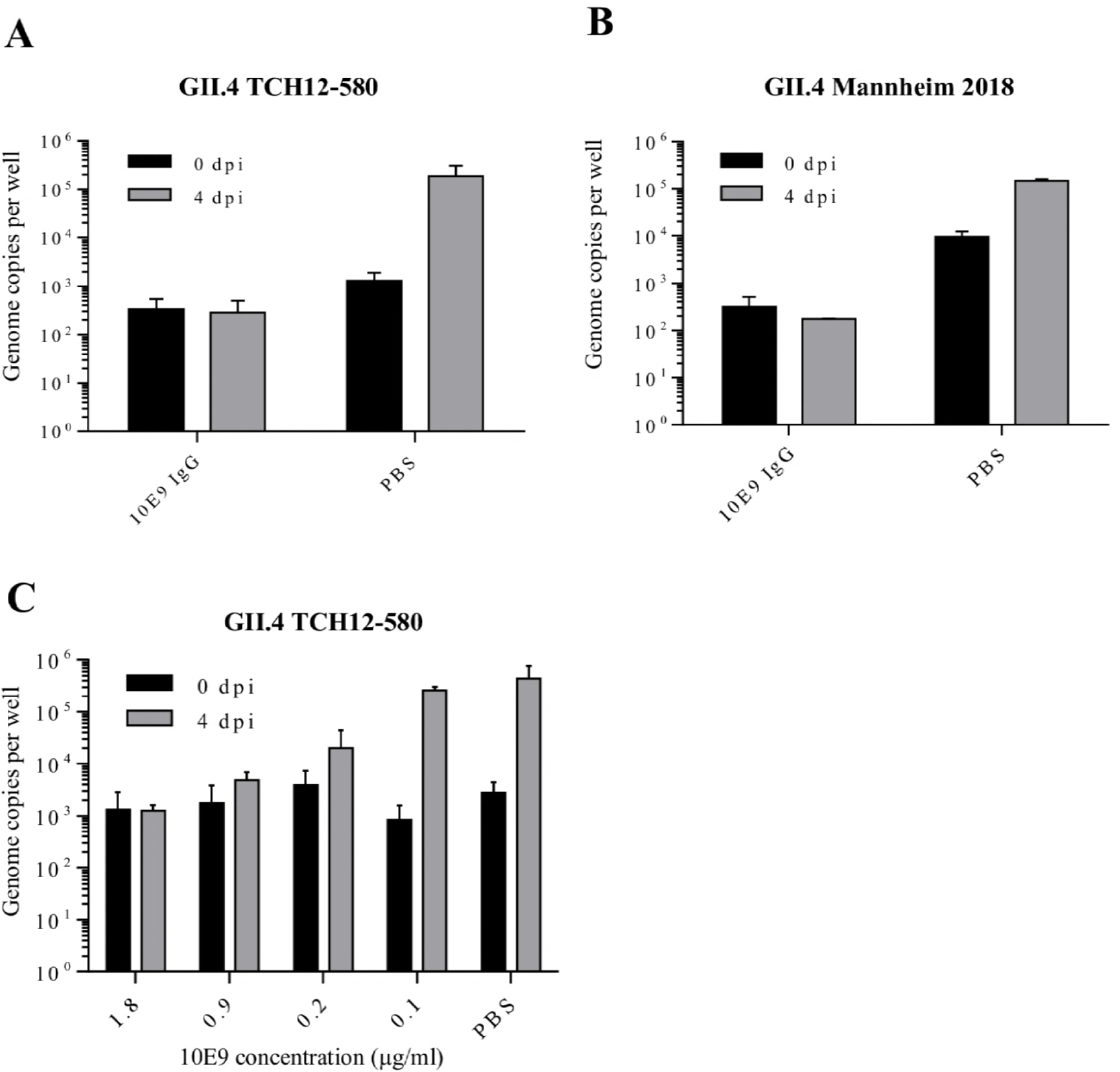
Inhibition of human norovirus replication by 10E9 IgG in human intestinal enteroid system. Neutralization properties of 10E9 IgG was tested in HIE using two GII.4 Sydney stool samples. GII.4 TCH12-580 sample (A) and GII.4 Mannheim 2018 (B) sample were completely neutralized by 10 μg/ml 10E9 IgG. (C) GII.4 Sydney 2012 TCH was pre-incubated with serially diluted 10E9 IgG prior to inoculation of jejunal monolayer. 10E9 MAb blocked norovirus replication with an IC_50_ value of 0.97 ng/ml. All experiments were performed in triplicates; standard deviation is shown with error bars.

## DISCUSSION

A plethora of studies have identified norovirus specific antibodies that block norovirus VLPs from binding to HBGAs (22-26). However, how these antibodies block the HBGA pocket is not completely understood. The mechanisms of HBGA binding inhibition can include sterical hindrance, allosteric inhibition and interference with capsid morphology (27, 43). Moreover, information on antigenic drift associated with the emergence on novel variants at the atomic level is lacking. In this current study, we were interested in explaining GII.4 antigenic drift and deciphering how amino acid substitutions might prevent antibody binding with closely related strains.

Our structural data, together with antibody binding and HBGA blocking results, indicated that the 10E9 MAb was GII.4 variant specific. The excessive substitutions of residues surrounding HBGA pocket could explain the lack of 10E9 MAb reactivity with older GII.4 strains, as the CDR regions of the antibody were tailored for the specific sequence of newer GII.4 variants. Interestingly, 10E9 Mab raised against GII.4 2006 strain could recognize and neutralize newer GII.4 2012 variant. Characterization of 10E9 MAb and its binding epitope is therefore especially relevant for the purpose of vaccine development against constantly evolving human norovirus. At the same time, the P domain residues interacting with the ABH-fucose moiety of HBGAs have for the most part remained highly conserved among CHDC-1974, Farmington Hills-2004, Saga-2006, and NSW-2012 GII.4 variants. Consistent with HBGA findings, another study of GII.4 variants also showed conserved HBGA binding profiles (16).

Antigenic changes in amino acid positions adjacent to the receptor binding site are frequently observed in another rapidly evolving RNA virus, influenza virus. In a striking difference to human norovirus, the temporal antigenic and genetic changes have caused the remarkable shifts in specificity and affinity for sialic acid receptors in pandemic influenza strains.

The mechanism of antigenic drift, while retaining HBGA binding may not be limited to the norovirus GII.4 variants, since another studies with GI specific human norovirus IgA antibody and GII.10 specific Nanobody displayed similar findings (23, 27). The structure of GI.1 specific IgA 5I2 showed that orientation of the bound Fab fragment was extraordinarily similar to that of 10E9 IgG. Both antibodies acted by steric interference and bound on the side of the HBGA pocket leaving the HBGA binding site still largely exposed. This mode of binding seems to be quite beneficial for the virus as it allows it to maintain a constant set of HBGA binding residues while modifying the surrounding region to escape antibody recognition. One hypothesis could be that the virus binds soluble HBGAs at the first encounter, which can than act as a shield for underlying residues, masking it from host antibody selection. On the other hand, since the receptor for the human norovirus remains unknown the binding site of neutralizing 10E9 MAb could potentially overlap with the yet unknown receptor site. Interestingly, both 10E9 Mab and 5I2 IgA Fabs clashed with the superimposed proteinaceous receptor for mouse norovirus CD300lf.

Overall, 10E9 antibody structural and functional characteristics illustrated that GII.4 antigenic drift might not interfere with HBGA binding. Our findings suggested that residues directly interacting with the HBGA maintain conservation, while flanking regions might experience antigenic variation that could result in the escape from neutralizing antibodies and lead to the emergence of novel antigenic variants. Ultimately, the temporal changes in antigenicity and amino acid variations in the surrounding HBGA pocket might reduce the ability of the host to defend against antigenic GII.4 variants, while retaining HBGA binding capabilities.

## ACKNOWLEDGEMENTS

The funding for this study was provided by the CHS foundation, the Helmholtz-Chinese Academy of Sciences (HCJRG-202), BMBF (Federal Ministry of Education and Research VIP+ 03VP00912), and DFG (FOR2327). We acknowledge the European Synchrotron Radiation Facility for provision of synchrotron radiation facilities and the Protein Crystallization Platform, CellNetworks, Heidelberg for assistance with protein crystallization. We also thank D. McAllister (ViroStat, USA) for providing the MAbs and 10E9 Fab sequence. We thank M. Estes, Baylor College of Medicine, Texas, USA and H. Schroten (Mannheim Children Hospital, Mannheim, Germany) for providing stool samples.

